# makeshift: a lightweight software for accessing and analyzing NMR data and protein dynamics

**DOI:** 10.64898/2026.08.17.745346

**Authors:** Gina El Nesr, Hannah K. Wayment-Steele

**Affiliations:** Biophysics Program, Stanford University; Department of Biochemistry, University of Wisconsin-Madison

## Abstract

Nuclear magnetic resonance (NMR) spectroscopy yields rich residue-level information on biomolecular dynamics and chemical environments, two frontiers for quantitative predictive methods in biochemistry. Decades of data are publicly archived in the Biological Magnetic Resonance Data Bank (BMRB)^1^, yet in practice, this information remains difficult to access and interpret at scale and within computational workflows. Here we present *makeshift*, an open-source Python package for accessing, curating, and analyzing NMR datasets. Users can readily retrieve and parse BMRB entries and perform essential analyses such as chemical shift re-referencing, secondary structure propensity prediction, and interpretation of relaxation datasets for dynamics. We re-implemented several widely-used NMR data calculations which were not open-source or available in Python and validated our implementations against the original implementations. By integrating data access, processing, and analysis into a single Python interface, *makeshift* lowers the barrier for reproducible, scalable analysis and machine learning applications using biomolecular NMR data.

## Introduction

Biomolecular dynamics is fundamental to understanding the functions of biomolecules, and NMR spectroscopy continues to serve as a gold standard in many contexts for how to measure even subtle conformational changes at ambient temperature in solution. Relaxation dispersion experiments are uniquely poised to measure populations and interconversion rates of otherwise “invisible” states. Furthermore, the central observable of NMR spectroscopy, the chemical shift, is a rich and poorly-predicted readout of chemical environments. Improved predictive power of chemical shifts would doubtlessly lead to models with improved understanding of chemical environments, with direct applications in domains such as drug discovery^2^ and enzyme engineering.

The Biological Magnetic Resonance Data Bank (BMRB)^1^ holds ~17,800 NMR macromolecular data entries as of August 2026, including NMR chemical shifts, relaxation constants, and order parameters. These measurements provide information that is often not captured by static structures, including residue-level conformational changes, flexibility, and dynamics across multiple timescales. Collectively, the BMRB is a valuable resource for comparative studies and machine learning on biomolecular structure and dynamics. In practice, however, most analyses using BMRB data rely on relatively small, manually-curated subsets of entries.

Differences in how chemical shift measurements and metadata are reported in the BMRB complicate comparisons between entries. Chemical shifts, for example, may contain uncorrected per-nucleus re-referencing offsets: Zhang et al. found that roughly 25% of entries with ^13^C assignments and 27% with ^15^N assignments needed significant re-referencing^3^. Residue numbering, relaxation units, and experimental conditions are also inconsistently reported. These differences should be standardized before combining, comparing, or using different entries.

Existing curation efforts and analysis methods address individual parts of this inconsistency problem but remain limited in scale or accessibility. RefDB^3^ was constructed in 2003 from a snapshot of the BMRB and has been intermittently updated, rendering other users dependent on external efforts inaccessible to themselves. Methods exist both to detect and correct referencing offsets and to derive or predict quantities that can be compared across entries, such as secondary-structure propensity or backbone flexibility. However, these tools are distributed separately as web servers, compiled binaries, or standalone scripts, each with their own input and output formats. Applying several methods across a data archive consequently requires substantial efforts to retrieve data, reconcile representations, convert formats, and connect outputs. By contrast, structural archives like the Protein Data Bank (PDB)^4^ helped enable breakthroughs like AlphaFold^5^ by making large datasets available in a consistent, readily processable format.

Here, we present makeshift, an open-source Python package for programmatic access to and curation of NMR datasets, with a focus on data related to protein dynamics (Figure 1a). makeshift retrieves publicly-available datasets and represents chemical shifts, sequences, experimental metadata, relaxation measurements, and related depositions through a consistent interface. It integrates tools for chemical shift re-referencing, spectrum and peak-list processing, secondary-structure analysis, and interpretation of relaxation datasets. makeshift also includes routines for processing newly-collected relaxation dispersion datasets. For methods intended to be reimplementations of existing algorithms, we validate each method against output from the original implementation and provide workflows and examples that can be applied to individual entries or archive-scale datasets. We anticipate that bringing together a suite of commonly-used NMR data analysis tools, with particular emphasis on observables related to dynamics at numerous timescales (Figure 1b), will enable users to readily develop software for using and interpreting NMR data. makeshift is freely available on GitHub, installable from PyPI, and documented at makeshift-docs.readthedocs.io.

**Figure 1.**
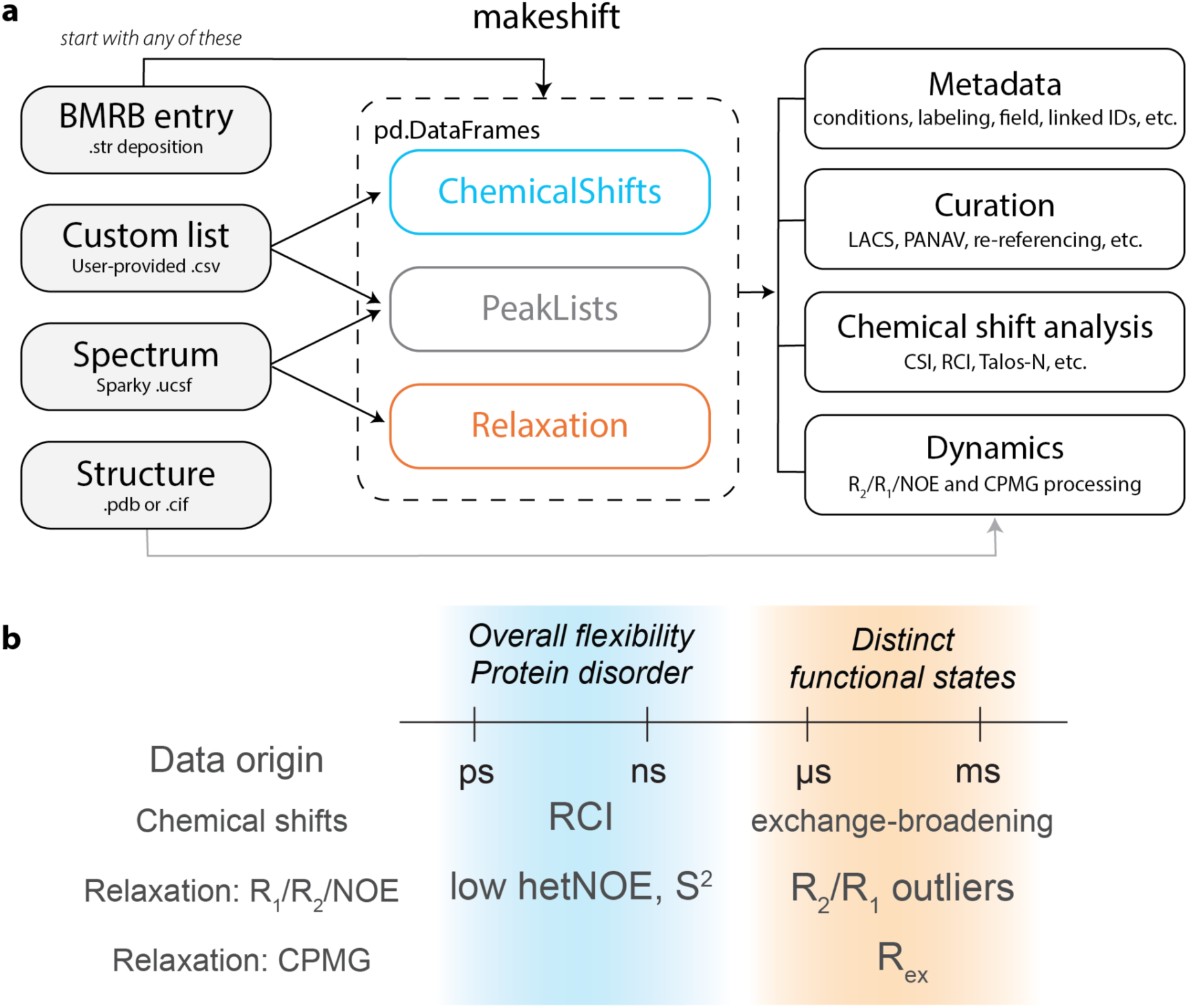
(a) Overview of makeshift inputs, data structures, and use-cases. (b) Overview of accessible data types in makeshift reporting on dynamics at different timescales.

### Development of makeshift

#### Data Fundamentals

##### Parsing and storing datasets

The core of publicly-available NMR data are chemical shift datasets. makeshift stores assigned chemical shifts (i.e. ChemicalShifts) into tables, with one row for each assigned atom. Each row contains the residue number, residue type, atom name, chemical shift, and reported uncertainty. Both BMRB and author residue numbering are retained because they may differ. The table also records which saveframe each assignment came from, keeping measurements under the same deposition but from different temperatures, conditions, or constructs separate.

Chemical shifts can also be organized as peak lists (i.e PeakLists), with one row per residue and one column per nucleus. makeshift standardizes alternative names for the same nucleus—for example, H, HN, and H1 for the backbone amide proton. Residues lacking the assignments required for a given analysis are omitted and recorded as being missing. Peak lists derived from BMRB assignments or imported from local files including other software formats can be analyzed the same way. Relaxation measurements follow a similar organization, with one row per residue and columns for the measured value, uncertainty, and source saveframes.

All tables use the widely-adopted pandas DataFrame format^6^, allowing users to inspect, filter, and analyze data with standard Python tools. The same analyses can also be applied to unpublished assignments or spectra, allowing users to compare their own data directly against archived entries (Figure 1a). makeshift standardizes differences in how data is reported and retains all metadata such as sample conditions, sample compositions, spectrometer, and linked database entries for easy access.

##### Chemical shift re-referencing

makeshift provides two re-referencing methods (LACS and PANAV) via the same set of commands (Figure 2a). These methods use different, complementary, underlying assumptions about data distributions and require no further inputs. LACS^7,8^ exploits the approximately linear relationship between an atom’s secondary chemical shift and the difference in secondary chemical shifts of the α- and β-carbons. We fit this relationship with a Huber regressor,from which we take the intercept as the nucleus offset (for N/HN, against the preceding residue’s difference in carbon shift). In comparison, PANAV^9^ first estimates each residue’s secondary structure from the H^α^ chemical shift, then calculates the referencing offset relative to the residue- and secondary-structure-specific shifts. Both methods report the estimated offset for each nucleus type and whether the estimate succeeded. Corrections are applied by adding the estimated offsets to the backbone chemical shifts. Figure 2b shows histograms of ^15^N and ^13^C shifts for BMRB entry 6586, which was first used in ref. ^9^ as a problematic entry; both PANAV and LACS recover consistent offsets in the same direction. Figure 2c compares offsets calculated in makeshift to those deposited in the BMRB of that atom type; we observe significant agreement with correlations generally well above 0.9. Other re-referencing methods exist but require data often inaccessible at scale. For example, SHIFTCOR^3,10^ was used to generate RefDB but requires predicted chemical shifts from a structure model.

**Figure 2.**
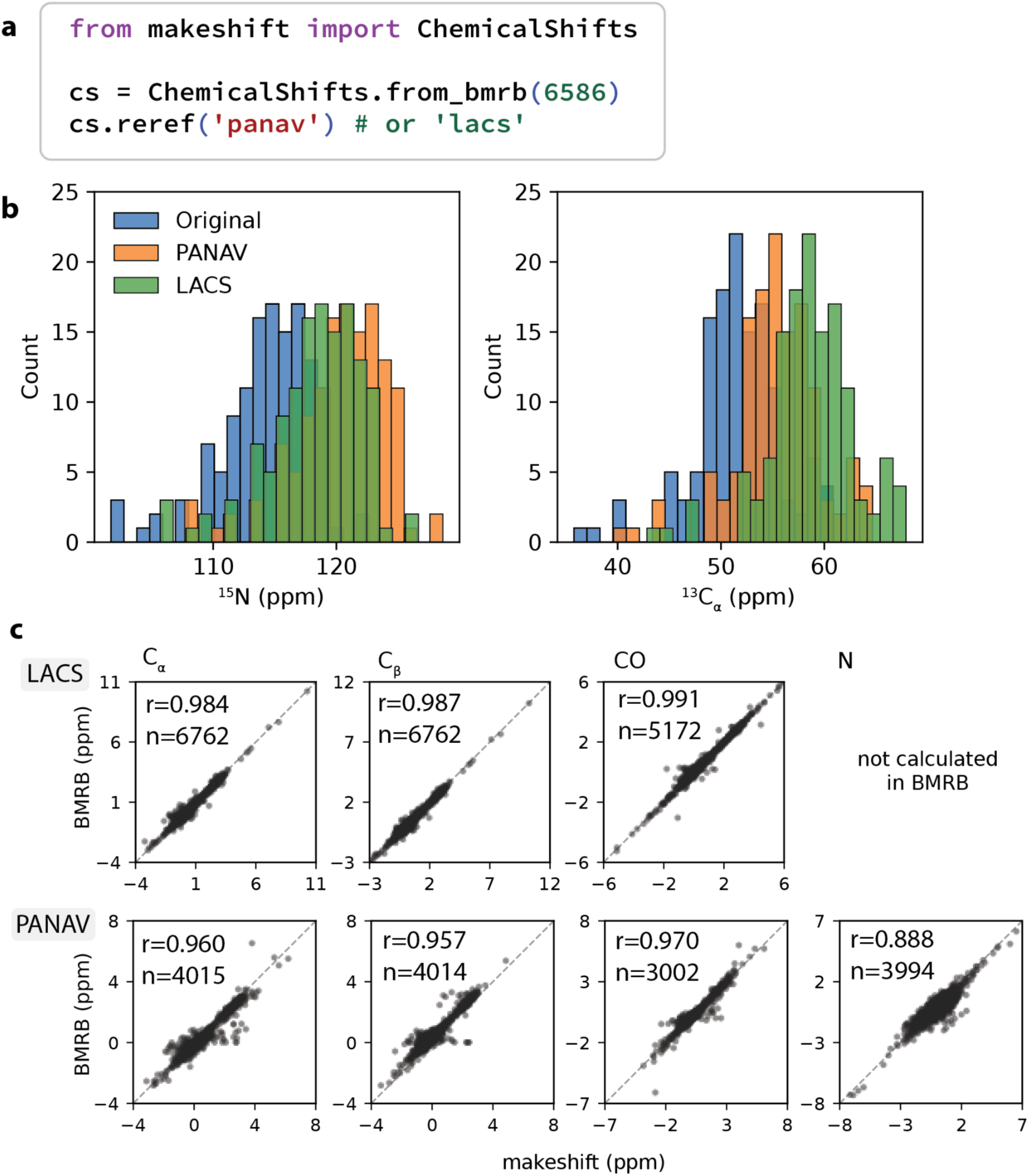
(a) Representative code to fetch a BMRB entry and run re-referencing to update the values in the dataset. (b) Distribution of data obtained for re-referenced BMRB entry 6586 for N shifts (left) and Ca shifts (right). (c) Comparison of makeshift LACS and PANAV re-implementations against available atom types in the BMRB.

#### Ambient solution structure: Chemical shift index and secondary-structure propensity

Chemical shifts have long been known to reflect structure properties such as secondary structure at room temperature. An established method for interpreting secondary structure is the chemical shift index (CSI)^11,12^, which we implement in makeshift. For each residue, both the difference from the expected random-coil chemical shift and a helix/sheet/coil classification are reported. We chose to implement the original CSI algorithms from Wishart et al, which classify each residue from secondary chemical shifts, treating glycine separately. For torsion angles, order parameters, and secondary structure predicted by a neural network, makeshift provides access to the TALOS-N software^13^.

#### Picosecond-nanosecond motions: Random coil index for backbone flexibility

Overall backbone disorder can be estimated using the random coil index (RCI)^14,15^, which derives a per-residue flexibility measure from the magnitude of chemical shift deviations from random coil values, smoothed along the sequence. Because RCI requires only assigned backbone shifts, it can be computed for any entry that contains them.

makeshift re-implements RCI in Python (representative code in Figure 3a) and provides two other backends. The first reproduces the reference script distributed with the original publication, while the second reproduces the implementation bundled within TALOS-N, which diverged from the original script. Both can be converted to estimated order parameters (S^2^ values) via an empirically-derived log transform^16^ (see Methods). Residues for which a value cannot be computed are reported as missing rather than 0. Example RCI data for the calmodulin-like domain of human non-muscle alpha-actinin 1 (BMRB 25871) is depicted in Figure 3b. Example RCI data for paired sets of folded and fully disordered / unfolded versions of the same construct are depicted in Figures 3c (apomyoglobin, BMRB entries 4061 and 16501, respectively) and Figure 3d (IscU, BMRB entries 17837 and 17836, respectively). Regions with RCI values above 0.1 typically correspond to regions with disorder.

**Figure 3.**
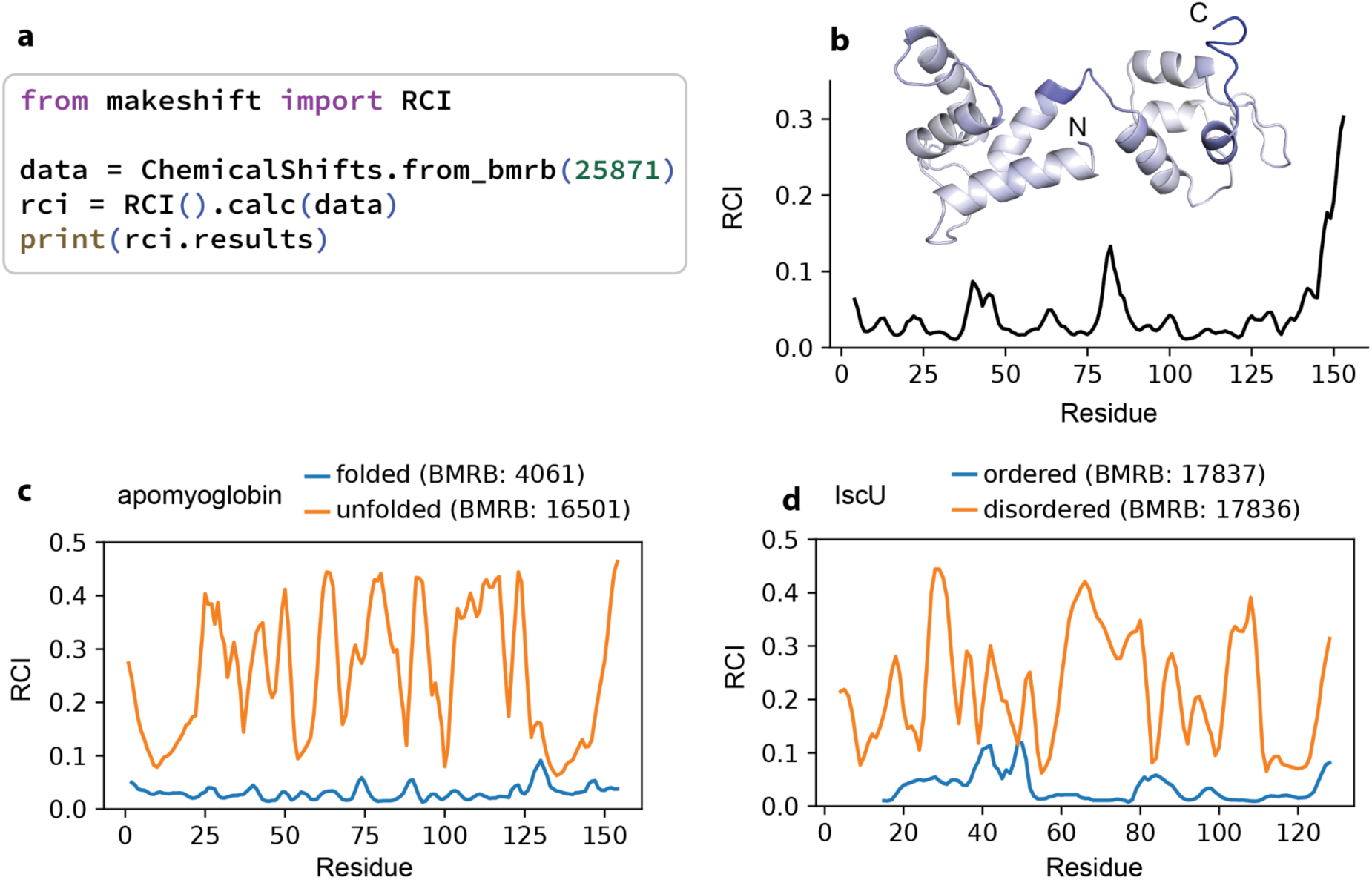
(a) Representative code to calculate random coil index (RCI) for a BMRB entry. (b) RCI values for the calmodulin-Like domain of human non-muscle alpha-actinin 1 (BMRB 25871). Inset: same RCI values visualized on structure (PDB: 2N8Z), dark blue = high RCI, light blue/white = low RCI. (c) RCI profiles for apomyoglobin in its folded, native state (BMRB 4061, blue) versus its acid-unfolded state at pH 2.4 (BMRB 16501, orange). (d) RCI profiles for IscU in its ordered (BMRB 17837, blue) versus disordered (BMRB 17836, orange) conformational states.

#### Micro-millisecond motions: NMR relaxation experiments

A frontier where NMR is uniquely poised in comparison to other data types is slow dynamics at the micro-to-millisecond (µs-ms) timescale, which relaxation experiments are specifically designed to measure. Since the early 90s, scientists have collected relaxation data on dynamics particularly relevant to biological function, with some deposited in the BMRB and others curated into datasets like RelaxDB^17^. To date, no journal deposition requirements exist for these datasets, which has hindered standardized curation and collection of experimental measurements of protein dynamics.

R_1_, R_2_, and hetNOE data are often interpreted using the model-free formalism^18,19^, but using this formalism has a number of limitations. Firstly, correctly fitting the standard model-free formalism requires relaxation data recorded at multiple static field strengths, which many deposited R_1_/R_2_/NOE datasets do not have (roughly 80% of datasets in RelaxDB contain a single field strength). Secondly, we have found in practice that fast-ModelFree^20^ is sensitive to the uncertainty estimates used, which do not follow a standard practice (e.g. sometimes estimated from signal/noise, sometimes estimated from duplicate points). The structure-based hydrodynamic calculation and labeling of residues with exchange implemented in makeshift (representative code in Figure 4a) avoids this requirement; given a structural model (experimental or predicted), residue-specific rigid-tumbling baseline is directly calculated. This allows for relaxation datasets to be compared and processed at scale, enabling the creation of datasets to be used in machine learning as demonstrated in ref. ^17^.

**Figure 4.**
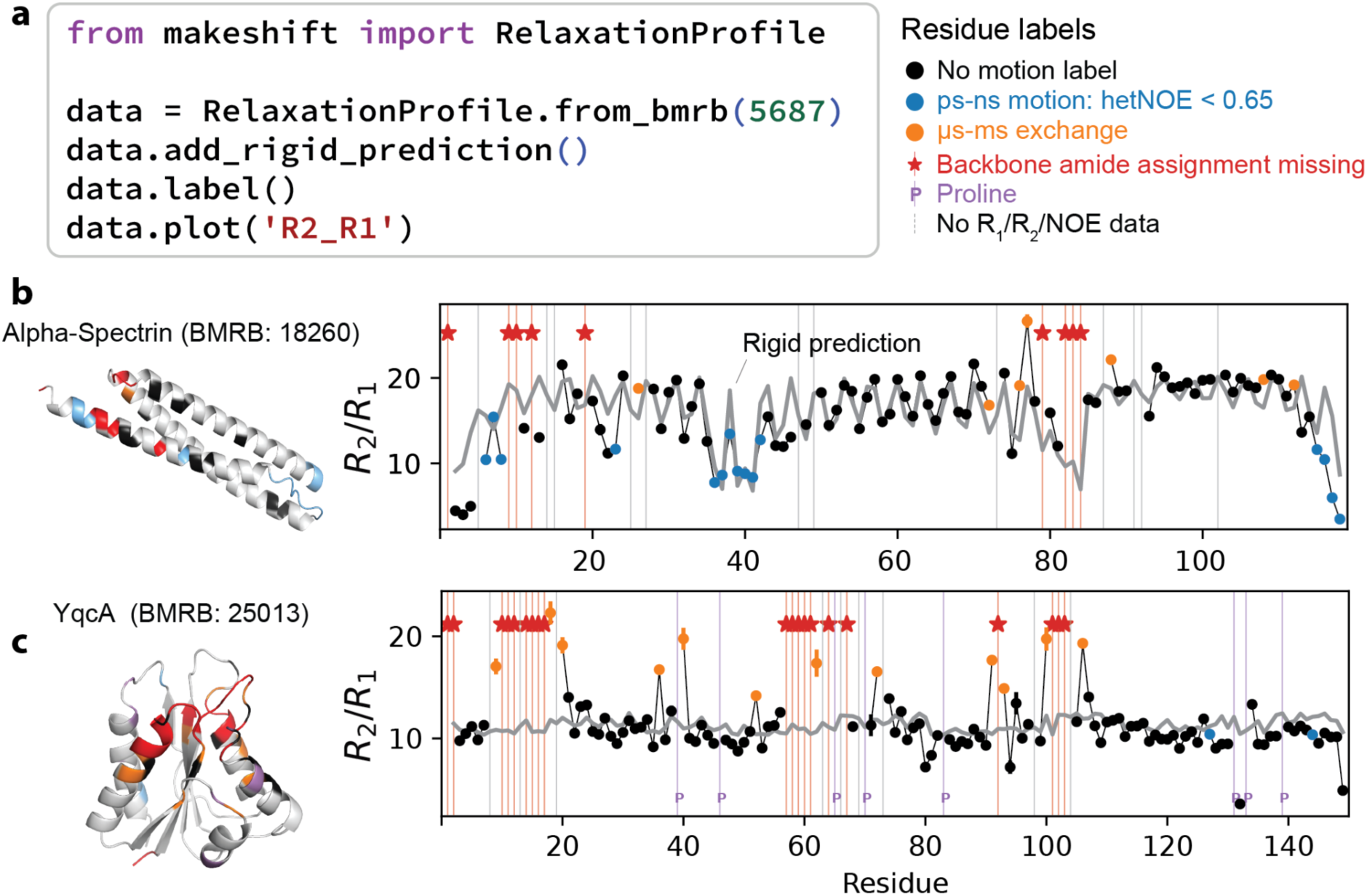
(a) Representative code to fetch relaxation data entry from BMRB and fit to quickly assess motions at different timescales from R_1_/R_2_/hetNOE data. (b) Representative data for protein alpha spectrin (BMRB entry 18260) with high anisotropic tumbling, seen in “zigzags” in rigid tumbling patterns. (c) Representative data for protein YqcA (BMRB entries 19151 and 25013 for assignments and parameters, respectively) with micro-to-millisecond exchange, seen in large spikes in R_2_/R_1_ over what rigid tumbling predicts.

In the widely-practiced R_1_/R_2_/hetNOE set of experiments, conformational exchange at a µs-ms timescale appears as excess R_2_/R_1_ above what a rigid, freely tumbling protein would show, so identifying it requires first predicting that baseline. For a rigid asymmetric rotor, the anisotropy of the rotational diffusion tensor causes the predicted R_2_/R_1_ ratio to vary between residues according to the orientation of each backbone N–H bond vector relative to the principal axes. Ignoring this variation can produce apparent exchange where relaxation is simply elevated by anisotropy, or mask genuine exchange where bond orientation is unfavorable. This distinction matters in practice: anisotropic tumbling alone can produce large, systematic per-residue variation in R_2_/R_1_ that is easily mistaken for exchange. In alpha-spectrin, for example (BMRB 18260), helical anisotropy produces a pronounced “zigzag” pattern in R_2_/R_1_ that tracks the alternating orientation of backbone N–H bonds along the helix (Figure 4b).

The processing and data-labeling pipeline used in ref.^17^ relies on HYDRONMR^21^ to calculate a rigid-tumbling baseline using shell-model hydrodynamics. HYDRONMR and its accelerated successor, Fast-HYDRONMR^22^, are distributed as compiled Fortran executables that require manually prepared input files, while the underlying source code is not openly available. These limitations make the software difficult to modify or run programmatically across hundreds of archived datasets. To make the complete workflow readily accessible and reproducible, we implemented an open-source version based on the double-sum approximation (DSA) introduced in Fast-HYDRONMR. The DSA recovers the same residue-level pattern of R_2_/R_1_ because the exact and approximate diffusion tensors share eigenvectors and have proportional eigenvalues.

makeshift returns the per-residue R_1_, R_2_ and R_2_/R_1_ from a given structure model. Because the intended application requires only a rigid-tumbling baseline for residue-level comparison against measured R_2_ or R_2_/R_1_, we focus on reproducing the relative profile rather than the absolute magnitude. As previously shown in ref. ^22^, the normalized deviation

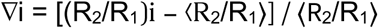

is largely insensitive to the atomic-element radius parameter and to whether the shell model or DSA is used, whereas the mean ⟨R_2_/R_1_⟩ contains a method- and parameter-dependent offset. We fit a single scaling constant that accounts for the difference between experimental overall tumbling time and that predicted by the HYDRONMR reimplementation. Residues whose experimental R_2_/R_1_ values substantially exceed the fitted rigid-tumbling baseline are then labeled to have µs-ms conformational exchange, as illustrated for apo YqcA flavodoxin protein (BMRB: 25013) in Figure 4c.

#### Processing and interpreting relaxation dispersion datasets

Carr-Purcell-Meiboom-Gill (CPMG) relaxation dispersion experiments^23,24^ provide a powerful means of characterizing dynamic processes on the µs-ms timescale. Going beyond R_1_/R_2_/hetNOE experiments, CPMG experiments can reveal otherwise invisible states and estimate their populations and rates of interconversion. CPMG relaxation dispersion experiments measure the effective transverse relaxation rate, R_2,eff_, as a function of the frequency of refocusing pulses applied during a fixed relaxation period, ν_CPMG_. Chemical exchange between states with distinct chemical shifts contributes additional relaxation, R_ex_, that adds to the intrinsic relaxation rate R_2_ and elevates R_2,eff_. The refocusing pulses interrupt this exchange-induced dephasing: as ν_CPMG_ increases, R_ex_ is progressively suppressed, and R_2,eff_ decreases toward a plateau. The shape of this dispersion profile encodes the exchange rate (k_ex_), the populations of the exchanging states, and the chemical shift difference between them (Δω). However, CPMG data must be collected at multiple magnetic field strengths or temperatures to determine the exchange regime and fit these parameters reliably^25^, and software to date focuses on these aspects of data fitting. There is an unmet need for software to quickly interpret simple markers of dynamics consistently across proteins starting from the raw data. When curating the RelaxDB-CPMG dataset^17^, we developed the following framework to process and interpret datasets from different proteins identically.

R_ex_ is typically estimated from the difference between R_2,eff_ measured at the lowest and highest CPMG field strengths. This calculation assumes that exchange broadening is fully suppressed at the highest field strength, such that the plateau approximates the exchange-free R_2_. For exchange occurring at the fast end of the µs-ms regime, however, even the highest experimentally-accessible CPMG field strengths may not fully suppress the exchange contribution. The plateau therefore remains elevated, causing the conventional difference-based estimate to underestimate, or entirely miss, this “unsuppressed” exchange. Including labels for these residues showed that Dyna-1 had greater predictive power than its original benchmark suggested^17^. Confidently identifying such residues requires an independent estimate of the exchange-free R_2_, which can be obtained from the same structure-based hydrodynamic calculation used for R_2_/R_1_, and scaled for each protein by a single fitted constant. makeshift automates this comparison as a standard step of its CPMG pipeline, flagging residues whose plateau R_2,eff_ exceeds the hydrodynamic prediction beyond experimental and fitting uncertainty, rather than requiring a bespoke, per-dataset reanalysis.

makeshift processes a CPMG relaxation-dispersion experiment as follows. Figure 5a depicts this processing pipeline on a representative dataset that makeshift automatically downloads as a demonstration for users: the NSH2 domain of SHP2 bound to the GAB-1 peptide^26^. makeshift picks peaks in the reference plane, assigns them against a reference peak list, fits per-plane lineshapes, calculates R_2,eff_(ν_CPMG_) = −(1/T)·ln(I/I_0_), and classifies each peak against a threshold derived from the hydrodynamic prediction (Figure 5a). makeshift bypasses the often manual process of mapping reference assignment data to dispersion data via a Hungarian algorithm. The resulting assignments can be visualized and adjusted in Python (Figure 5b). Uncertainties are propagated from the lineshape fits and, where duplicate CPMG field strength (ν_CPMG_) points are present, from the deviation between them. Resulting R_2,eff_ values and uncertainties can be visualized in conventional plots (Figure 5c) as well as in a protein-wide “fountain plot” (Figure 5d). Finally, the processed data can be exported and fit to determine parameters like p_B_, k_ex_, and dω in other programs such as ChemEx^27^.

**Figure 5.**
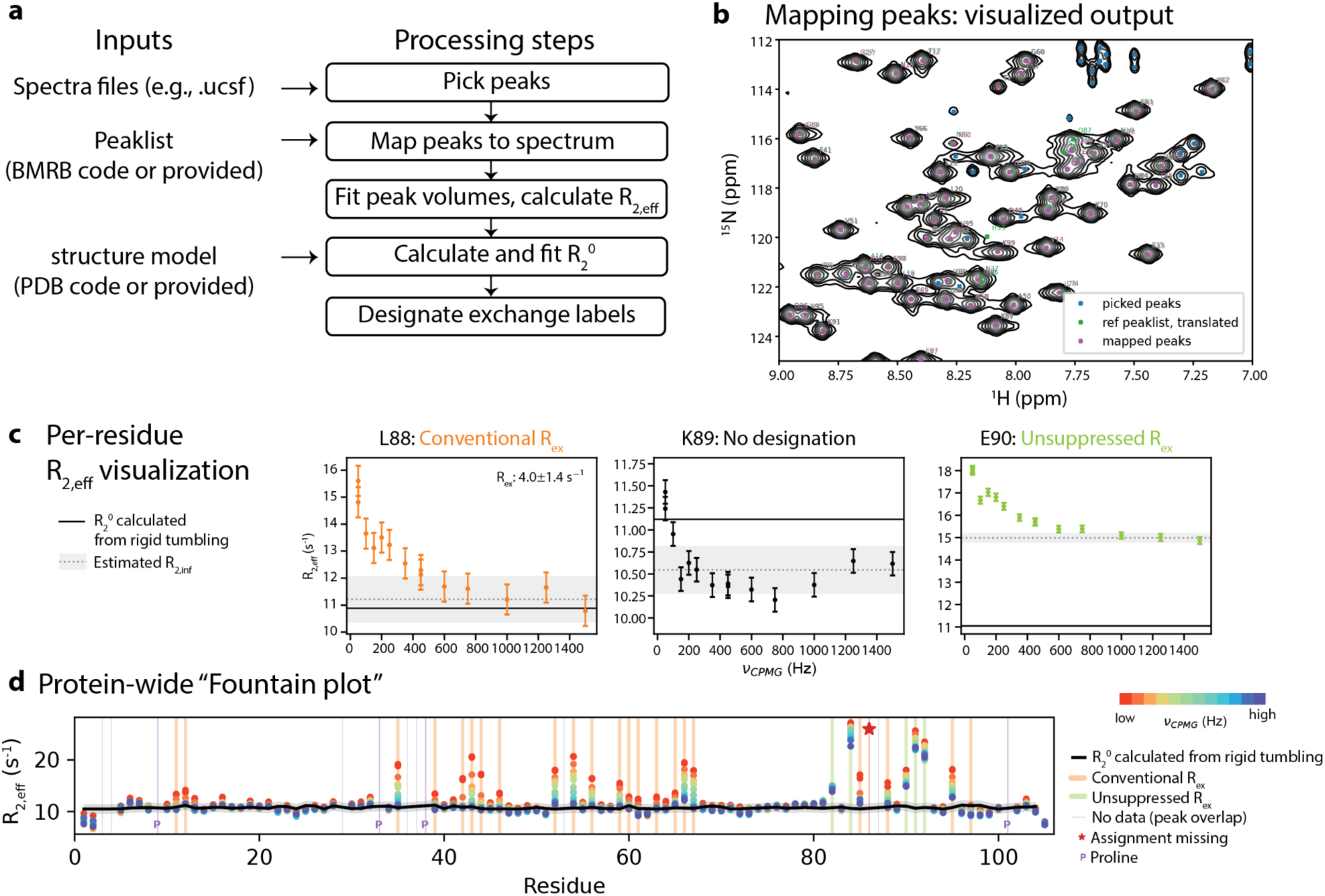
(a) Pipeline in makeshift for processing user-provided CPMG data. (b) Visualizing output of automated peak-mapping routine. (c) Visualizing R_2,eff_ per residue. (d) Visualizing global view of exchange across the entire protein.

## Methods

### Validation of makeshift

We validated each reimplemented method against outputs from the original implementation.

#### Re-referencing with LACS and PANAV

makeshift implements two structure-independent chemical shift re-referencing methods: LACS and PANAV. We validated each method against the corresponding BMRB-hosted reference: deposited LACS.str offsets for LACS and the BMRB entry validate API for PANAV. For every usable macromolecule entry for which a LACS.str file exists, the chemical shifts and offsets were calculated and paired against the BMRB reference. Agreement is summarized as Pearson r and RMSD over all paired entries (Table 1). We note that the BMRB does not deposit an offset for ^15^N and HN for LACS, so LACS N is not compared.

**Table 1.** Validation of makeshift LACS/PANAV offsets against BMRB reference values.

| Method | Atom | n | Pearson r | RMSD (ppm) |
| --- | --- | --- | --- | --- |
| LACS | CA | 6762 | 0.984 | 0.146 |
| LACS | CB | 6762 | 0.987 | 0.136 |
| LACS | CO | 5172 | 0.991 | 0.107 |
| PANAV | N | 3994 | 0.888 | 0.401 |
| PANAV | CA | 4015 | 0.960 | 0.276 |
| PANAV | CB | 4014 | 0.957 | 0.406 |
| PANAV | CO | 3002 | 0.970 | 0.192 |

LACS agrees closely with the deposited references for all three ^13^C nuclei. PANAV recovers the BMRB validate offsets well for the carbon nuclei, but with larger residual scatter than LACS, as expected for a secondary-structure dependent, multi-atom protocol. Agreement is weaker for N.

#### Random Coil Index

makeshift’s Random Coil Index (RCI) predictor supports two interchangeable backends. The default, algorithm=“wishart”, is a direct port of the reference implementation^14^ file rci_v_1c.py. We validated it against the exact input/output pair distributed with the reference script (BMRB entry 4403, the J domain of murine polyomavirus T antigen), reproducing the reference RCI values to floating-point precision (Pearson r = 1.0, RMSD = 7.9 x 10^-14^, maximum absolute deviation = 4.1 x 10^-13^ across n = 79 residues).

A second backend, algorithm=“talosn”, ports the related but algorithmically distinct RCI-derived order-parameter (S^2^) calculation bundled inside TALOS-N^13^. No independent reference test case exists for this module, so instead we validated it by running the compiled TALOS-N binary directly and comparing its per-residue S^2^ output against makeshift’s port across nine BMRB entries chosen for diversity in size and chemical-shift completeness (856 residues total; Table 2). Pooled across all nine entries, the two agree with Pearson r = 0.989 (median absolute S^2^ deviation 0.0010; 82.1% of residues within 0.01 and 96.0% within 0.05 of the reference value). The residual disagreement is concentrated at proline and glycine residues and traces to a TALOS-N homology-search fallback (calcAverageCS) not ported.

**Table 2.** Per-entry validation of algorithm=“talosn” against the compiled TALOS-N binary.

| BMRB | n | Pearson r | median $\vert \Delta\vert $ | max $\vert \Delta\vert $ |
| --- | --- | --- | --- | --- |
| 11080 | 97 | 0.9955 | 0.0009 | 0.0829 |
| 15451 | 89 | 0.9952 | 0.0009 | 0.0679 |
| 15490 | 58 | 0.9877 | 0.0018 | 0.0689 |
| 15521 | 67 | 0.9640 | 0.0004 | 0.1412 |
| 15581 | 140 | 0.9979 | 0.0008 | 0.0598 |
| 15763 | 67 | 0.7654 | 0.0084 | 0.1040 |
| 15959 | 108 | 0.8278 | 0.0004 | 0.3209 |
| 52018 | 90 | 0.9919 | 0.0009 | 0.1050 |
| 5991 | 140 | 0.9877 | 0.0010 | 0.0949 |
| **Pooled** | **856** | **0.989** | **0.0010** | — |

#### HydroNMR calculation

makeshift.hydronmr predicts per-residue backbone ^15^N T_1_, T_2_, and NOE relaxation from a static PDB structure, reimplementing the core hydrodynamic and spectral-density calculations of Fast-HYDRONMR. We validated it against Fast-HYDRONMR output for seven proteins spanning 158–266 modeled residues in RelaxDB-CPMG dataset from ref. ^17^ (AQADK, BLAC, BLVRB, CHI19, CYPA, KRAS, VHR).

makeshift’s current hydrodynamic bead model approximates each heavy atom as a uniformly sized sphere, rather than the calibrated, residue-specific shell model used by the original Fast-HYDRONMR runs; this introduces a systematic scale offset in absolute R_2_/R_1_ values but is expected to preserve the relative, per-residue pattern of mobility. We therefore compared mean-subtracted per-residue R_2_/R_1_ (each residue’s R_2_/R_1_ relative to its structure’s own mean) between implementations. Agreement was high across all seven structures (Pearson r = 0.925–0.9996; Table 3), with the lowest correlation (KRAS, r = 0.925) still capturing the dominant features of the per-residue mobility profile.

**Table 3.**
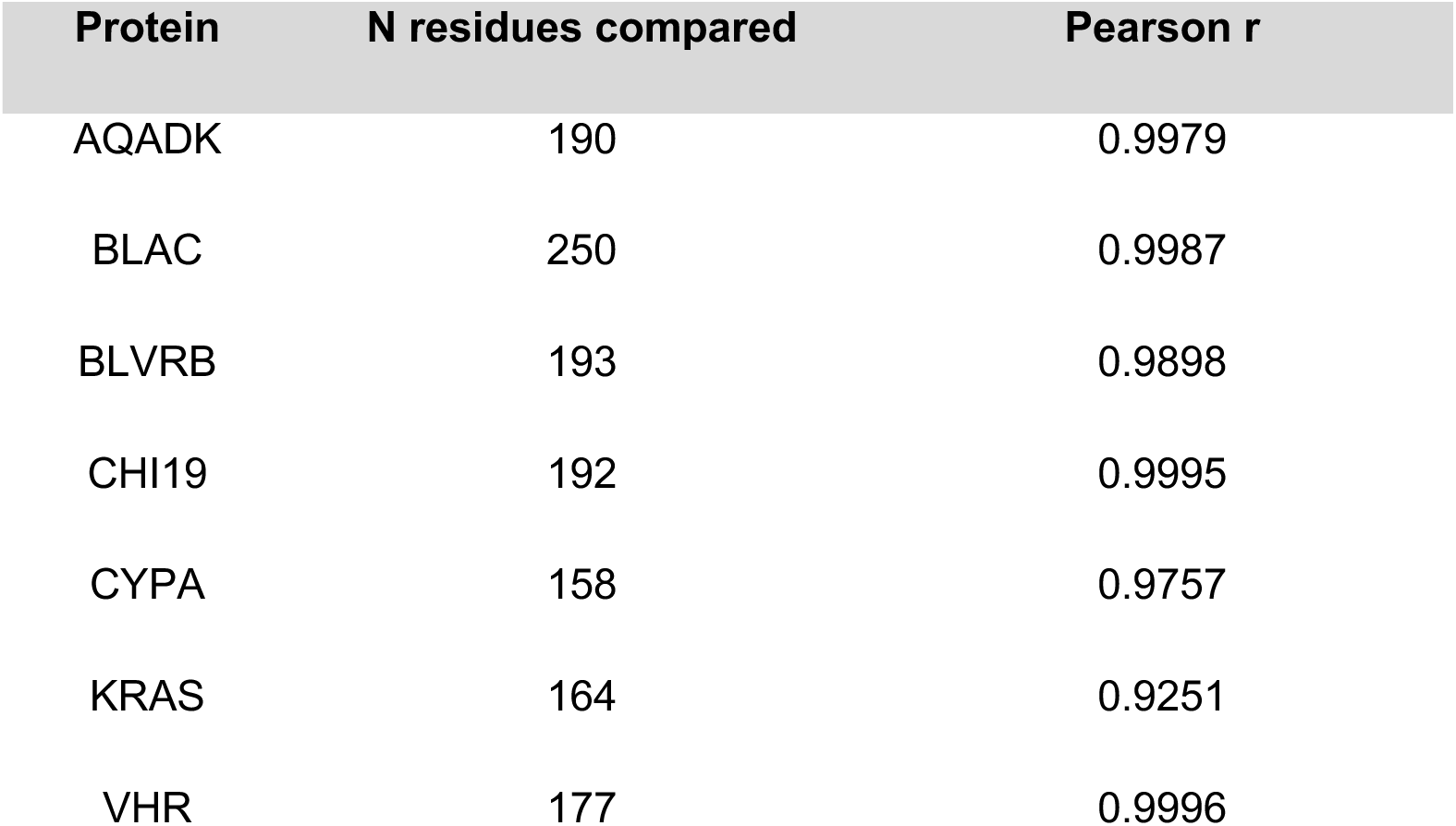
Per-protein validation of makeshift.hydronmr against Fast-HYDRONMR (mean-subtracted R_2_/R_1_) using reference AlphaFold2 models of proteins in RelaxDB-CPMG.

| Protein | N residues compared | Pearson r |
| --- | --- | --- |
| AQADK | 190 | 0.9979 |
| BLAC | 250 | 0.9987 |
| BLVRB | 193 | 0.9898 |
| CHI19 | 192 | 0.9995 |
| CYPA | 158 | 0.9757 |
| KRAS | 164 | 0.9251 |
| VHR | 177 | 0.9996 |

## Discussion

Rapid advancements in predictive structural biology have been largely enabled by centralized, machine-readable archives like the Protein Data Bank. Biomolecular NMR provides complementary, residue-level insight into local chemical environments and in solution dynamics; yet existing workflows for NMR data curation rely heavily on standalone web servers, compiled executables, and non-standardized formats. By bringing data retrieval, curation, and analysis into a lightweight and open-source Python package, makeshift lowers the barrier for large-scale datasets for biomolecular structure and dynamics.

We provide makeshift as a package that integrates simple, interpretable, and widely-used processing methods in NMR rather than exhaustively reimplementing advanced predictors. Each of these methods is validated against the original implementation where one exists. We also provide a programmatic framework to access and process dynamics datasets at scale; makeshift automates hydrodynamic baseline calculations and labeling of relaxation dispersion datasets (R_1_/R_2_/hetNOE and CPMG) to enable standardized curation of dynamics corpora like RelaxDB.

By bridging decades’ worth of NMR archives with modern computational frameworks, we believe that makeshift will facilitate access to rich datasets for training predictive models targeting protein dynamics, enzyme engineering, and drug discovery.

## Data and code availability

makeshift is distributed under an MIT license and is available on GitHub at https://github.com/WaymentSteeleLab/makeshift and on PyPI as makeshift-nmr. Documentation is made available at https://makeshift-docs.readthedocs.io/.

## Acknowledgements

We thank the Biological Magnetic Resonance Bank for maintaining and sharing NMR data, and the Bax laboratory at the NIH for sharing details on TALOS-N implementation. We thank Nevin Pai, Cizhang Zhao, and Leonel Bustamante-Carballo for beta-testing makeshift and providing valuable feedback. AI tools were used to prepare some parts of software and manuscript text, with outputs reviewed, modified, and confirmed by authors.

## Accession codes

Data from BMRB entries 6586, 18260, 19151, 25013, 25871, 4061, 16501, 17836, and 17837 were used for figures in this manuscript.

## Notes

https://github.com/WaymentSteeleLab/makeshift/tree/main

https://makeshift-docs.readthedocs.io/

### Competing Interest Statement

The authors have declared no competing interest.

## References

1. Hoch, J. C. et al. Biological Magnetic Resonance Data Bank. Nucleic Acids Res. 51, D368– D376 (2023).

2. Pellecchia, M. et al. Perspectives on NMR in drug discovery: a technique comes of age. Nat. Rev. Drug Discov. 7, 738–745 (2008).

3. Zhang, H., Neal, S. & Wishart, D. S. RefDB: a database of uniformly referenced protein chemical shifts. J Biomol NMR 25, 173–195 (2003).

4. Berman, H. M. et al. The Protein Data Bank. Nucleic Acids Res. 28, 235–242 (2000).

5. Jumper, J. et al. Highly accurate protein structure prediction with AlphaFold. Nature 596, 583–589 (2021).

6. Team, T. P. D. Pandas-Dev/pandas: Pandas. (Zenodo, 2020). doi:10.5281/zenodo.3509134.

7. Wang, L., Eghbalnia, H. R., Bahrami, A. & Markley, J. L. Linear analysis of carbon-13 chemical shift differences and its application to the detection and correction of errors in referencing and spin system identifications. J. Biomol. NMR 32, 13–22 (2005).

8. Wang, L. & Markley, J. L. Empirical correlation between protein backbone 15N and 13C secondary chemical shifts and its application to nitrogen chemical shift re-referencing. J. Biomol. NMR 44, 95–99 (2009).

9. Wang, B., Wang, Y. & Wishart, D. S. A probabilistic approach for validating protein NMR chemical shift assignments. J. Biomol. NMR 47, 85–99 (2010).

10. Neal, S., Nip, A. M., Zhang, H. & Wishart, D. S. Rapid and accurate calculation of protein 1H, 13C and 15N chemical shifts. J. Biomol. NMR 26, 215–240 (2003).

11. Wishart, D. S., Sykes, B. D. & Richards, F. M. The chemical shift index: a fast and simple method for the assignment of protein secondary structure through NMR spectroscopy. Biochemistry 31, 1647–1651 (1992).

12. Wishart, D. S. & Sykes, B. D. The 13C chemical-shift index: a simple method for the identification of protein secondary structure using 13C chemical-shift data. J. Biomol. NMR 4, 171–180 (1994).

13. Shen, Y. & Bax, A. Protein backbone and sidechain torsion angles predicted from NMR chemical shifts using artificial neural networks. J. Biomol. NMR 56, 227–241 (2013).

14. Berjanskii, M. V. & Wishart, D. S. A simple method to predict protein flexibility using secondary chemical shifts. J. Am. Chem. Soc. 127, 14970–14971 (2005).

15. Berjanskii, M. V. & Wishart, D. S. Application of the random coil index to studying protein flexibility. J. Biomol. NMR 40, 31–48 (2008).

16. Berjanskii, M. V. & Wishart, D. S. The RCI server: rapid and accurate calculation of protein flexibility using chemical shifts. Nucleic Acids Res. 35, W531–7 (2007).

17. Wayment-Steele, H. K. et al. Learning millisecond protein dynamics from what is missing in NMR spectra. Nature 1–3 (2026) doi:10.1038/s41586-026-10989-4.

18. Lipari, G. & Szabo, A. Model-free approach to the interpretation of nuclear magnetic resonance relaxation in macromolecules. 1. Theory and range of validity. J. Am. Chem. Soc. 104, 4546–4559 (1982).

19. Lipari, G. & Szabo, A. Model-free approach to the interpretation of nuclear magnetic resonance relaxation in macromolecules. 2. Analysis of experimental results. J. Am. Chem. Soc. 104, 4559–4570 (1982).

20. Cole, R. & Loria, J. P. FAST-Modelfree: a program for rapid automated analysis of solution NMR spin-relaxation data. J. Biomol. NMR 26, 203–213 (2003).

21. García de la Torre, J., Huertas, M. L. & Carrasco, B. HYDRONMR: prediction of NMR relaxation of globular proteins from atomic-level structures and hydrodynamic calculations. J. Magn. Reson. 147, 138–146 (2000).

22. Ortega, A. & García de la Torre, J. Efficient, accurate calculation of rotational diffusion and NMR relaxation of globular proteins from atomic-level structures and approximate hydrodynamic calculations. J. Am. Chem. Soc. 127, 12764–12765 (2005).

23. Chiliveri, S. C. & Deshmukh, M. V. Recent excitements in protein NMR: Large proteins and biologically relevant dynamics. J. Biosci. 41, 787–803 (2016).

24. Hansen, D. F., Vallurupalli, P. & Kay, L. E. Using relaxation dispersion NMR spectroscopy to determine structures of excited, invisible protein states. J. Biomol. NMR 41, 113–120 (2008).

25. Sauerwein, A. C. & Hansen, D. F. Relaxation Dispersion NMR Spectroscopy. in Protein NMR 75–132 (Springer US, Boston, MA, 2015). doi:10.1007/978-1-4899-7621-5_3.

26. Glaser, A. W., Pádua, R. A. P., Ojoawo, A. M., Sullivan, C. & Kern, D. Phosphatase SHP2 pathogenic mutations enhance activity by altering conformational sampling. Proc. Natl. Acad. Sci. U. S. A. 123, e2513851123 (2026).

27. Tharatipyakul, A., Numnark, S., Wichadakul, D. & Ingsriswang, S. ChemEx: information extraction system for chemical data curation. BMC Bioinformatics 13 Suppl 17, S9 (2012).

