## Supporting Information for "makeshift: a lightweight software for accessing and analyzing NMR data and protein dynamics"

### HydroNMR Processing Methods

#### *Overview of the pipeline*

Given a protein structure in PDB format, makeshift.hydronmr proceeds through the following stages: (1) construction of a primary hydrodynamic bead model from the atomic coordinates; (2) assembly and inversion of the grand mobility matrix to obtain the translational, rotational, and coupling friction tensors; (3) construction of the $6 \times 6$ generalized diffusion tensor and extraction of the $3 \times 3$ rotational diffusion tensor $D_{rr}$; (4) eigendecomposition of $D_{rr}$ to obtain principal values and the five Woessner correlation times; (5) estimation of backbone amide N–H bond vectors from PDB coordinates; and (6) per-residue computation of the anisotropic spectral density and NMR relaxation rates. All calculations are implemented in pure Python using NumPy for array operations and SciPy for LAPACK access.

#### *Bead model construction*

The primary hydrodynamic model is constructed in the AER (Atomic Element Radius) mode, following references (García de la Torre et al. 2000; García De La Torre et al. 2000; Carrasco and García de la Torre 1999). Each non-hydrogen atom in the PDB file is replaced by a spherical bead of radius *a* = 3.0Å centered at the atomic coordinates, giving a model with *N* beads where *N* is the number of non-hydrogen atoms. PDB ATOM and HETATM records are parsed using fixed-column offsets per the PDB format specification. The default AER of 3.0Å is the consensus value from the original HYDRONMR (García de la Torre et al. 2000) validation, representing the van der Waals radius of a typical heavy atom plus approximately one monolayer of hydration. All coordinates are converted to centimeters for cgs-unit consistency throughout.

#### *Grand mobility matrix and RPY hydrodynamic interaction*

The $3N \times 3N$ grand mobility matrix $B$ is assembled following the standard bead-model formalism (Garcia de la Torre and Bloomfield 1981; Carrasco and García de la Torre 1999). Diagonal $3 \times 3$ blocks are $(1/a_i)I$, corresponding to the Stokes friction of an isolated sphere. Off-diagonal $3 \times 3$ blocks $T_{ij}$ are given by the Rotne–Prager–Yamakawa (RPY) hydrodynamic interaction tensor (Rotne and Prager 1969; Yamakawa 1970):

$$T_{ij} = \frac{6}{8r_{ij}}\{[1 + a_{ij}^2]\, I + [1 - 3a_{ij}^2]\hat{r}_{ij} \otimes \hat{r}_{ij}\}$$

where $r_{ij} = |r_j - r_i|$ is the inter-bead separation,

$$a_{ij}^2 = \frac{a_i^2 + a_j^2}{3r_{ij}^2}, \qquad \hat{r}_{ij} = \frac{r_j - r_i}{r_{ij}}$$

is the unit separation vector. The full matrix is multiplied by $1/(6\pi\eta)$, where η is the solvent viscosity (default 0.01 poise). The current implementation uses the non-overlap RPY branch; all pairwise computations are vectorized using NumPy broadcasting, giving $O(N^2)$ array operations.

#### *Inversion to friction tensors*

The grand mobility matrix $B$ is inverted to give the grand friction matrix $C = B^{-1}$ via LAPACK packed-symmetric Cholesky factorization and inversion (DPPTRF / DPPTRI), accessed through SciPy's LAPACK wrapper, following the approach of García de la Torre *et al.* (2000). The $3 \times 3$ translational ($C_t$), rotational ($C_r$), and translation–rotation coupling ($C^c$) friction tensors are obtained by summing over the $3 \times 3$ block structure of $C$ about a chosen origin:

$$C_t = \sum_{ij} C_{ij}, \qquad C^c = \sum_{ij} S_i C_{ij}, \qquad C_r = \sum_{ij} S_i C_{ij} S_j^T$$

where $S_i$ is the $3 \times 3$ skew-symmetric matrix of $(r_i - r_0)$. The center of diffusion — the origin that minimizes rotational–translational coupling — is found by solving

$$Ad = b, \qquad A = Tr(C_t)I - C_t,$$

where $b$ encodes the antisymmetric part of $C^c$ (García de la Torre and Bloomfield, 1977). The friction tensors are then recomputed about this center.

#### *Generalized diffusion tensor*

The $6 \times 6$ generalized friction supermatrix is assembled as

$$\Xi = [C_t\ (C^c)^T\ C^c\ C_r\ ]$$

and the generalized diffusion tensor is

$$D = k_B T\, \Xi^{-1},$$

where $k_B = 1.380649 \times 10^{-16}\ erg\ K^{-1}$ and $T$ is the absolute temperature (default 293 K). The $3 \times 3$ rotational diffusion tensor $D_{rr}$ is the lower-right block of $D$. All quantities are in cgs units.

#### *Rotational diffusion tensor eigendecomposition and correlation times*

The symmetric $3 \times 3$ rotational diffusion tensor $D_{rr}$ is diagonalized via numpy.linalg.eigh, giving principal values $D_x \geq D_y \geq D_z$ and their corresponding eigenvectors. The mean rotational diffusion coefficient is

$$D_r = \frac{D_x + D_y + D_z}{3},$$

with anisotropy

$$\Delta = \left(D_x^2 + D_y^2 + D_z^2 - D_x D_y - D_y D_z - D_z D_x\right)^{1/2}.$$

The five Woessner correlation times are:

$$\tau_0 = \frac{1}{6D_r - 2\Delta}, \qquad \tau_1 = \frac{1}{4D_y + D_x + D_z}, \qquad \tau_2 = \frac{1}{4D_x + D_y + D_z},$$

$$\tau_3 = \frac{1}{4D_z + D_x + D_y}, \qquad \tau_4 = \frac{1}{6D_r + 2\Delta}.$$

The harmonic mean correlation time

$$\tau_c = \frac{1}{6D_r}$$

is also returned; this is the single isotropic correlation time relevant for orientation-averaged calculations (García De La Torre et al. 2000).

#### *N–H bond vector estimation*

For each residue, the backbone amide N–H bond unit vector is estimated from PDB heavy-atom coordinates, without requiring explicit hydrogen positions. The standard sp² geometry at the backbone nitrogen is exploited: because N, $C'_{i-1}$, $C\alpha_i$ , and are approximately coplanar at ~120° bond angles, the N–H direction is opposite the bisector of the $C'_{i-1}$–N–$C\alpha_i$ angle:

$$n = -normalize[normalize(N - C'_{i-1}) + normalize(N - C\alpha_i)].$$

This is the standard implicit-hydrogen approximation used by HYDRONMR (Iflag = 1). Proline residues and the N-terminal residue of each chain lack either a preceding C′ or an amide hydrogen and are skipped. For a typical 150-residue protein this produces vectors for approximately 140 residues, consistent with Fast-HYDRONMR (Ortega and García de la Torre 2005) output.

#### *Anisotropic spectral density and NMR relaxation rates*

Per-residue NMR relaxation rates are computed using the full five-term anisotropic spectral density function of (Woessner 1962). Given the direction cosines $l_1, l_2, l_3$ of the N–H unit vector along $D_x, D_y, D_z$ respectively, the Woessner amplitudes are:

$$A_1 = 3l_1^2 l_3^2, \qquad A_2 = 3l_2^2 l_3^2, \qquad A_3 = 3l_1^2 l_2^2,$$

$$f' = l_1^4 + l_2^4 + l_3^4 - \frac{1}{3},$$

$$g_\delta = (l_1^4 + 2l_2^2 l_3^2)D_x + (l_2^4 + 2l_1^2 l_3^2)D_y + (2l_1^2 l_2^2 + l_3^4)D_z,$$

$$A_0 = \frac{3}{4}\left(f' + \frac{g_\delta - D_r}{\Delta}\right), \qquad A_4 = \frac{3}{4}\left(f' - \frac{g_\delta - D_r}{\Delta}\right).$$

The amplitudes satisfy

$$\sum_{k=0}^{4} A_k = 1.$$

The spectral density evaluated at frequency $\omega$ is:

$$J(\omega) = \sum_{k=0}^{4} A_k \frac{2}{5} \frac{\tau_k}{1 + \omega^2 \tau_k^2}.$$

Longitudinal and transverse relaxation rates and the NOE enhancement are given by the standard dipolar–CSA expressions (García De La Torre et al. 2000; Kay et al. 1989):

$$R_1 = \frac{d^2}{4}[J(\omega_H - \omega_N) + 3J(\omega_N) + 6J(\omega_H + \omega_N)] + c^2 J(\omega_N),$$

$$R_2 = \frac{d^2}{8}[4J(0) + J(\omega_H - \omega_N) + 3J(\omega_N) + 6J(\omega_H) + 6J(\omega_H + \omega_N)] + \frac{c^2}{6}[4J(0) + 3J(\omega_N)],$$

$$NOE = 1 + \frac{d^2}{4R_1}\frac{\gamma_H}{\gamma_N}[6J(\omega_H + \omega_N) - J(\omega_H - \omega_N)],$$

where

$$\omega_H = \gamma_H B_0, \qquad \omega_N = \gamma_N B_0,$$

are the $^{1}$H and $^{15}$N Larmor frequencies:

$$d^2 = \left(\frac{\mu_0}{4\pi}\right)^2 \frac{\hbar^2 \gamma_H^2 \gamma_N^2}{r_{NH}^6}$$

is the squared dipolar coupling constant; and

$$c^2 = \frac{\omega_N^2 \Delta\sigma^2}{3}$$

is the squared CSA contribution. Default parameters match the ground-truth validation cases: $B_0 = 11.74\ T$ (500 MHz ¹H), $\gamma_H = 2.675 \times 10^8\ rad\ s^{-1}\ T^{-1}$, $\gamma_N = -2.7126 \times 10^7\ rad\ s^{-1}\ T^{-1}$, $r_{NH} = 1.02$ Å, and $\Delta\sigma = -172\ ppm$.

***Limitations***

Our implementation uses the double-sum approximation (DSA) directly on the primary bead model, without the shell-model extrapolation to zero bead size that the original HYDRONMR employs. As described by Ortega and García de la Torre (2005), the DSA introduces a scalar scale factor into $D_{rr}$ — identical for all three eigenvalues — so the absolute value of $\langle T_1/T_2 \rangle$ is offset from experiment, but the relative anisotropy $\bar{V}_i$ is correctly reproduced. For applications that require accurate absolute correlation times (e.g., validation against $^{75}$Se or $^{15}$N $R_2$ magnitudes), we recommend the original Fast-HYDRONMR program.

The current implementation is also memory-limited for very large proteins: the $3N \times 3N$ dense mobility matrix requires $O(N^2)$ memory. Structures with more than approximately 5,000 non-hydrogen atoms will approach or exceed available RAM. Future work could implement the shell-model approach (tiling each atom's surface with minibeads and extrapolating across shell sizes), which both improves accuracy and reduces the effective *N* by replacing per-atom beads with a much smaller number of surface minibeads.

### References


Carrasco, B., and J. García de la Torre. 1999. “Hydrodynamic Properties of Rigid Particles: Comparison of Different Modeling and Computational Procedures.” *Biophysical Journal* 76 (6): 3044–3057.

Garcia de la Torre, J. G., and V. A. Bloomfield. 1981. “Hydrodynamic Properties of Complex, Rigid, Biological Macromolecules: Theory and Applications.” *Quarterly Reviews of Biophysics* 14 (1): 81–139.

García De La Torre, J., M. L. Huertas, and B. Carrasco. 2000. “Calculation of Hydrodynamic Properties of Globular Proteins from Their Atomic-Level Structure.” *Biophysical Journal* 78 (2): 719–730.

García de la Torre, J., M. L. Huertas, and B. Carrasco. 2000. “HYDRONMR: Prediction of NMR Relaxation of Globular Proteins from Atomic-Level Structures and Hydrodynamic Calculations.” *Journal of Magnetic Resonance (San Diego, Calif.: 1997)* 147 (1): 138–146.

Kay, L. E., D. A. Torchia, and A. Bax. 1989. “Backbone Dynamics of Proteins as Studied by 15N Inverse Detected Heteronuclear NMR Spectroscopy: Application to Staphylococcal Nuclease.” *Biochemistry* 28 (23): 8972–8979.

Ortega, Alvaro, and Jose García de la Torre. 2005. “Efficient, Accurate Calculation of Rotational

Diffusion and NMR Relaxation of Globular Proteins from Atomic-Level Structures and Approximate Hydrodynamic Calculations.” *Journal of the American Chemical Society* 127 (37): 12764–12765.

Rotne, Jens, and Stephen Prager. 1969. “Variational Treatment of Hydrodynamic Interaction in Polymers.” *The Journal of Chemical Physics* 50 (11): 4831–4837.

Woessner, D. E. 1962. “Nuclear Spin Relaxation in Ellipsoids Undergoing Rotational Brownian Motion.” *The Journal of Chemical Physics* 37 (3): 647–654.

Yamakawa, Hiromi. 1970. “Transport Properties of Polymer Chains in Dilute Solution: Hydrodynamic Interaction.” *The Journal of Chemical Physics* 53 (1): 436–443.
